# Testicular but not ovarian hormones shape patch-leaving adaptation and impulsive choice in rats

**DOI:** 10.64898/2026.07.08.737251

**Authors:** Yadong Dai, Kaylyn Castillo, James R. Hinman

## Abstract

Hormonal regulation of patch-leaving decision-making remains poorly understood. Here, young adult male and female Long-Evans rats were tested in a patch-leaving task before and after orchiectomy (ORCH), ovariectomy (OVX), or sham surgery, and were subsequently assessed in an impulsive-choice task. Patch leaving was measured under long– and short-travel conditions. Before surgery, males showed longer overstay than females during long-travel sessions, whereas no clear sex difference was detected during short-travel sessions. After surgery, orchiectomy did not produce a uniform shift in patch leaving but selectively disrupted the progressive reduction in overstay that normally emerged across repeated long-travel sessions. By contrast, ovariectomy produced weaker effects and did not reveal a comparably robust change in female patch leaving. Spatial and idle occupancy analyses showed that gonadectomy also altered within-patch behavior, with orchiectomy most strongly increasing idling-related measures in males, whereas ovariectomy more strongly redistributed female patch occupancy. Estrous stage did not significantly organize pre-surgical female overstay. Greater impulsive choice was associated with smaller post-surgical reductions in long-travel overstay in the unadjusted analysis. Together, these findings indicate that testicular hormones selectively support patch-leaving adaptation under high travel cost.

## Introduction

Animals often forage in environments where resources are distributed unevenly across space and decline as they are exploited, requiring decisions about whether to continue harvesting a current patch or leave to search elsewhere ^1–3^. Optimal foraging theory proposes that animals should allocate behavior to maximize net resource intake under constraints such as travel cost, patch quality, and opportunity cost ^1,2^. The marginal value theorem predicts that a forager should leave a depleting patch when the instantaneous rate of return in the current patch falls below the average rate of return available in the environment ^1,2^. This framework has been useful for explaining patch-leaving behavior across species, including rodents, non-human primates, and humans ^4–11^.

Laboratory patch-foraging paradigms are especially valuable because they preserve key ecological features of foraging while allowing experimental control over reward depletion, travel cost, and task history ^3^. In rodents, patch residence time increases when travel cost is high and decreases when reward availability declines, indicating that animals integrate multiple sources of environmental information during stay-or-leave decisions ^3–6,10–12^. However, rodent and human foragers often remain in patches longer than predicted by reward-maximizing models, a pattern described as overharvesting or overstaying ^4,6,8–11^. This tendency suggests that patch leaving reflects more than simple reward-rate maximization and may incorporate uncertainty, learning, energetic cost, opportunity cost, and other motivational variables^7–11^.

Sex is one biological factor that may contribute to individual differences in patch-leaving behavior. In a spatial patch-foraging task, female rats leave patches earlier than males and show less overharvesting, while locomotor differences explain only part of this sex effect ^10,11^. Gonadal hormones are plausible mechanisms for such sex-dependent variation because estradiol and testosterone influence neural systems involved in reward valuation, action selection, motivation, and learning ^13–15^. In female rodents, ovarian hormones regulate memory-system engagement and alter effort-based decision-making and reward-seeking ^13,14,16–18^. In males, androgen signaling affects mesocorticolimbic and executive-function circuits relevant to cost-benefit decisions, and testosterone manipulations influence impulsivity-related behavior ^15,19^. Although these studies do not directly address patch leaving, they provide a mechanistic basis for testing whether the removal of testes or ovaries might alter sequential foraging decisions.

Patch leaving may also be related to impulsive choice. Delay-discounting tasks measure preference between smaller immediate rewards and larger delayed rewards, with a stronger preference for the immediate option interpreted as a demonstration of greater impulsive choice behavior ^20–23^. Although patch foraging and delay discounting differ procedurally, both require animals to evaluate current reward availability against delayed future benefits based on the sense of time ^4,20,23^. In a depleting patch, staying provides immediate but declining reward opportunities, whereas leaving imposes a travel delay before access is gained to another patch ^4,7,10,11^. Consistent with this conceptual overlap, rats show similar biases in patch-foraging and intertemporal-choice tasks, suggesting that these behaviors may depend on partly shared computations^4^. Neural systems implicated in intertemporal choice, including frontal cortical, anterior cingulate, and broader corticolimbic valuation circuits, are also involved in foraging and cost-benefit decision making ^2,5,7,22,23^.

Sex differences in impulsive choice have been reported in humans and rodents, although their direction and magnitude vary across species, strain, age, reward conditions, and task design ^20,24–27^. On a delay discounting task, female rats show a greater preference for smaller immediate rewards than males, with estrous stage not significantly modulating this pattern ^20^. Critically, ovariectomy did not alter impulsive choice in females, while orchiectomy increased impulsive choice in males, suggesting that testicular hormones help maintain male-typical tolerance for delayed reward ^20^. This finding, together with evidence that testosterone administration reduces impulsivity-related behavior in male rats, suggests that testicular hormones may regulate delay– or persistence-related decision processes ^19,20^.

Despite growing interest in naturalistic decision-making models, two gaps remain. First, although sex differences in patch-leaving behavior have been reported, no study has directly tested whether gonadectomy alters patch residence time or overstaying in rats ^10,11^. Second, although patch leaving and delay discounting share an intertemporal structure and may recruit overlapping decision processes, the relationship between individual differences in patch-leaving performance and impulsive choice has not been directly examined in the same animals ^4,20^. Addressing these gaps may clarify whether sex differences in patch-leaving foraging are mediated by activational effects of gonadal hormones and whether patch-leaving reflects a broader impulsive-choice phenotype.

The present study addressed these questions in adult male and female Long-Evans rats tested on a patch-leaving task before and after gonadectomy or sham surgery. Animals first completed patch-leaving sessions under short– and long-travel conditions. Males then received either orchiectomy or sham surgery, and females received either ovariectomy or sham surgery. Following recovery, all animals were retested on the same patch-foraging task. Estrous cycle stage was monitored in females during both the pre-surgical and post-surgical patch-leaving phases. All rats subsequently completed a delay-discounting task. Based on prior evidence that males overharvest more than females in patch-foraging tasks and that orchiectomy increases impulsive choice, we hypothesized that removal of testicular hormones would reduce overstaying, particularly in long-travel sessions, in orchiectomized males relative to sham males ^10,11,20^. In contrast, based on evidence that ovariectomy does not alter impulsive choice in female rats, we predicted that ovariectomy would produce little or no change in patch-leaving behavior relative to sham females ^20^. Finally, we tested whether individual differences in patch-leaving performance covaried with impulsive choice, which would support the possibility that sequential foraging and intertemporal choice share common decision-making mechanisms ^4,10,11,20,23^.

## Results

### Baseline sex differences in patch-leaving behavior before gonadectomy

Young adult Long-Evans rats (N = 24 males, N = 24 females) were tested between approximately postnatal day 70 and 120. All rats first completed a pre-surgical patch-leaving task consisting of 30-min sessions under the same reward-rate schedule for 10 days. During this baseline phase, rats completed five sessions with a long corridor travel interval (60 s), followed by five sessions with a short corridor travel interval (5 s). All patches used the same depleting food-dispensing rate, which was refreshed upon each new patch entry. After baseline testing, rats of each sex were assigned to either gonadectomy or sham surgery. Males received either orchiectomy or sham surgery, whereas females received either ovariectomy or sham surgery. Following a two-week recovery period, all animals were tested again on the same patch-leaving protocol used before surgery (Fig. 1A).

**Figure 1.**
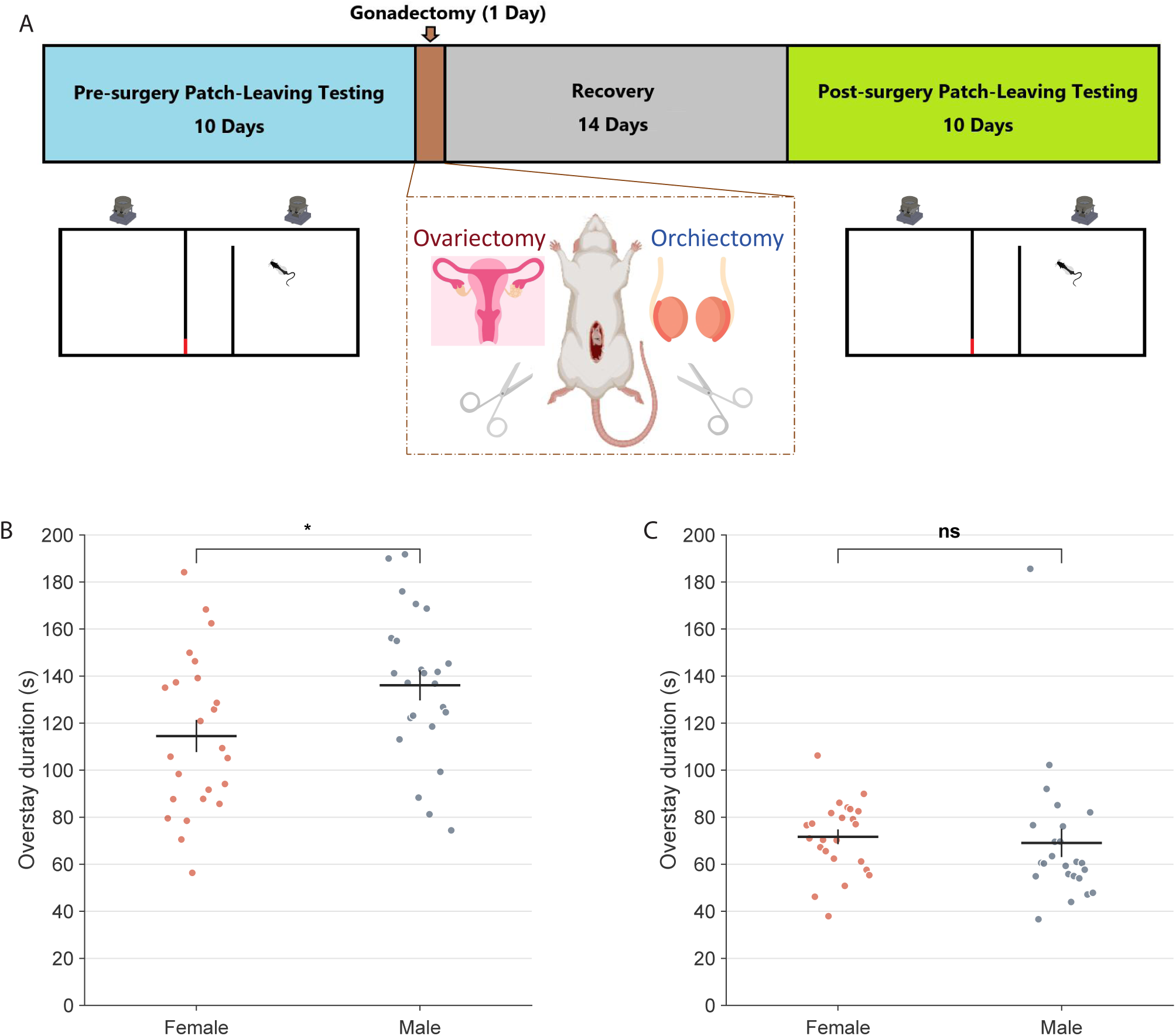
Experimental design and baseline sex differences in patch-leaving overstay. A, schematic of the experimental sequence, including pre-surgical patch-leaving testing, gonadectomy or sham surgery, recovery, and post-surgical patch-leaving retesting. B, rat-level mean overstay duration during pre-surgical long-travel sessions. C, rat-level mean overstay duration during pre-surgical short-travel sessions. In panels B and C, each point represents one rat averaged across the five pre-surgical sessions of that travel condition; horizontal bars indicate means and vertical bars indicate SEM. *p < 0.05; ns, not significant.

We first examined whether male and female rats differed in baseline patch-leaving behavior before surgical manipulation. During pre-surgical long-travel sessions, male rats showed longer overstay durations than female rats, indicating that males remained in depleting patches longer before leaving when patch switching was costly (Fig. 1B; males: 136.08 ± 6.41 s; females: 114.52 ± 6.83 s; Wilcoxon rank-sum test, p = 0.0296). In contrast, no significant sex difference was detected during pre-surgical short-travel sessions (Fig. 1C; males: 69.05 ± 5.97 s; females: 71.66 ± 3.13 s; Wilcoxon rank-sum test, p = 0.0969). Thus, baseline sex differences in patch-leaving were selective to the long-travel condition and were not evident when travel cost was low.

### Orchiectomy selectively attenuates late-stage overstay reduction during long-travel patch leaving in males

We next examined whether gonadectomy altered post-surgical patch-leaving behavior in males. When long-travel data were pooled across the five sessions, both orchiectomized and sham males showed lower post-surgical overstay durations than before surgery (Fig. 2A; orchiectomy: 133.35 ± 8.99 s pre, 97.99 ± 7.57 s post; sham: 142.41 ± 9.60 s pre, 102.56 ± 9.27 s post). However, rat-level mixed-effects analysis indicated that the male long-travel effect was not best described as a uniform pre– to post-surgical shift. Instead, orchiectomy altered the session-dependent trajectory of post-surgical performance, such that overstay declined across days in sham males but was markedly blunted in orchiectomized males (post-surgical surgery × session interaction: F(1,110) = 4.21, p = 0.0424). This divergence was most apparent in the final long-travel session, when day-matched post-surgical overstay reduction (pre-surgical minus post-surgical overstay) remained robust in sham males but was near zero or negative in orchiectomized males (Fig. 2E; sham: 40.86 ± 16.53 s; orchiectomy: –9.23 ± 9.41 s; Wilcoxon rank-sum test, p = 0.0035).

**Figure 2.**
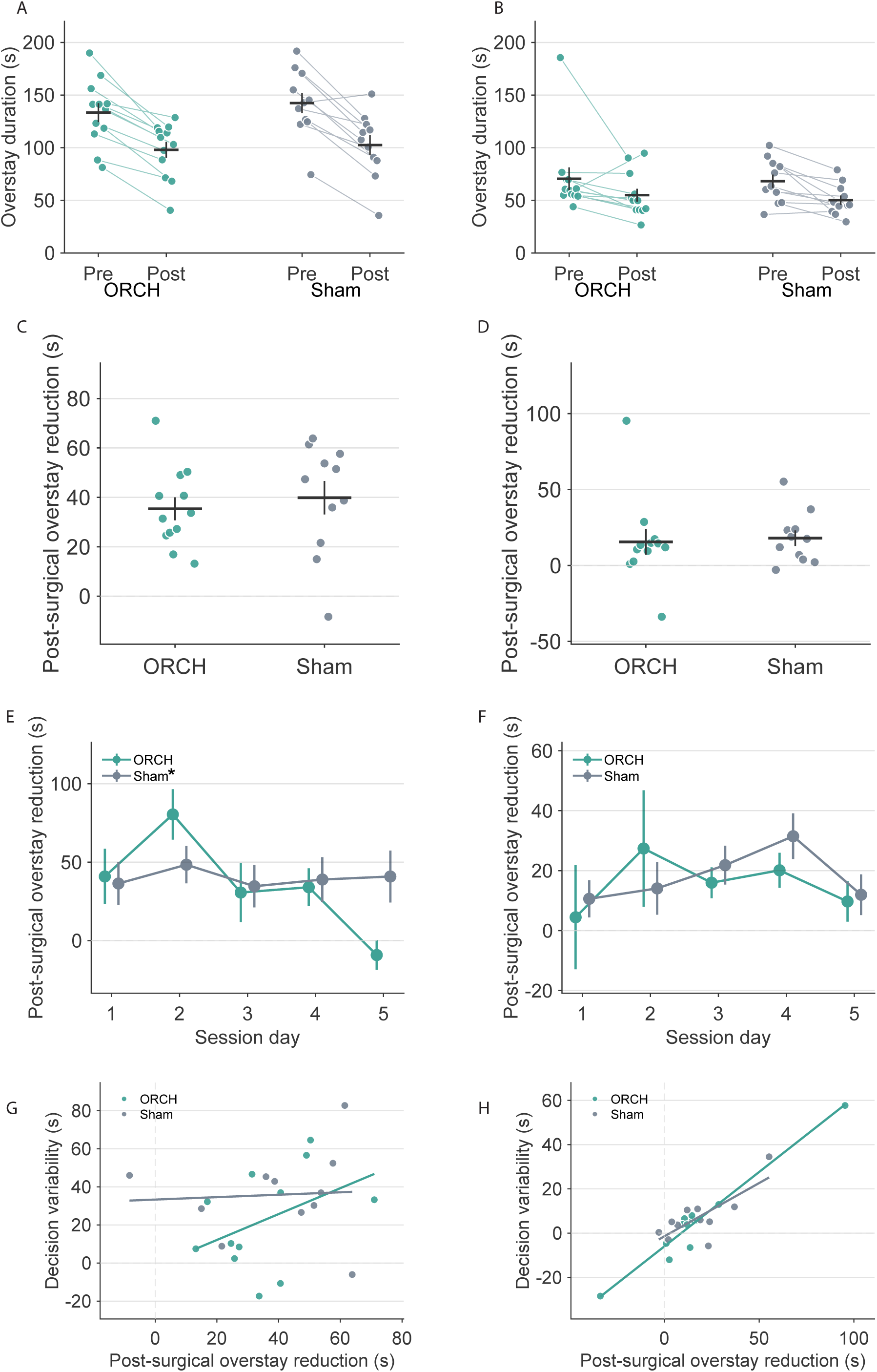
Orchiectomy selectively alters long-travel overstay adaptation in males. A, pooled pre-surgical and post-surgical mean overstay duration in long-travel sessions for ORCH and Sham males. B, pooled pre-surgical and post-surgical mean overstay duration in short-travel sessions. C, pooled long-travel post-surgical overstay reduction. D, pooled short-travel post-surgical overstay reduction. E, day-matched long-travel post-surgical overstay reduction across the five test sessions. F, day-matched short-travel post-surgical overstay reduction across the five test sessions. G, relationship between long-travel post-surgical overstay reduction and decision variability. H, relationship between short-travel post-surgical overstay reduction and decision variability. In panels A-D and G-H, each point represents one rat. In panels E and F, points and error bars indicate means ± SEM. For panels C-H, post-surgical overstay reduction was calculated as pre-surgical minus post-surgical overstay duration, such that larger positive values indicate a larger decrease in overstay after surgery. In panel E, the asterisk beside Sham in the legend denotes a significant overall surgery-by-session interaction in the male post-surgical long-travel trajectory.

By contrast, short-travel behavior did not show an analogous surgery-dependent reorganization. Although pooled short-travel overstay was lower after surgery in both male groups (Fig. 2B; orchiectomy: 70.50 ± 10.76 s pre, 54.97 ± 6.09 s post; sham: 68.24 ± 6.26 s pre, 50.25 ± 4.39 s post), the post-surgical short-travel trajectory did not differ between orchiectomized and sham males (post-surgical surgery × session interaction: F(1,110) = 0.0023, p = 0.9617). Accordingly, pooled post-surgical overstay reduction did not differ significantly between male groups in either long-travel or short-travel sessions (Fig. 2C, D; long: p = 0.3722; short: p = 0.5181).

We also asked whether individual differences in post-surgical overstay reduction were related to within-rat overstay variability. In males, long-travel overstay reduction was not significantly associated with long-travel overstay variability (Fig. 2G; Spearman rho = 0.302, p = 0.1606). A positive association was observed in the short-travel condition (Fig. 2H; rho = 0.734, p = 0.0001), but because orchiectomy did not alter short-travel adaptation across days, the clearest behavioral consequence of testicular hormone removal was a selective disruption of the progressive reduction in overstay that normally emerged across repeated long-travel experience.

### Female long-travel overstay showed a weaker post-surgical change, and pre-surgical overstay was not detectably organized by estrous stage

We next performed the corresponding analysis in females. In the pooled long-travel data, sham females showed lower post-surgical overstay than before surgery, whereas ovariectomized females showed a smaller and less consistent reduction (Fig. 3A; sham: 121.04 ± 8.91 s pre, 83.20 ± 6.27 s post; ovariectomy: 110.59 ± 11.01 s pre, 89.05 ± 13.09 s post). At the rat level, however, the post-surgical long-travel trajectory did not differ significantly between ovariectomized and sham females, although the effect trended in that direction (post-surgical surgery × session interaction: F(1,110) = 3.66, p = 0.0584). Likewise, the overall pre– versus post-surgical term did not reach significance in the female long-travel model (F(1,230) = 3.51, p = 0.0624). Thus, unlike the male dataset, female long-travel behavior did not show a clearly resolved surgery-dependent change across post-surgical sessions.

**Figure 3.**
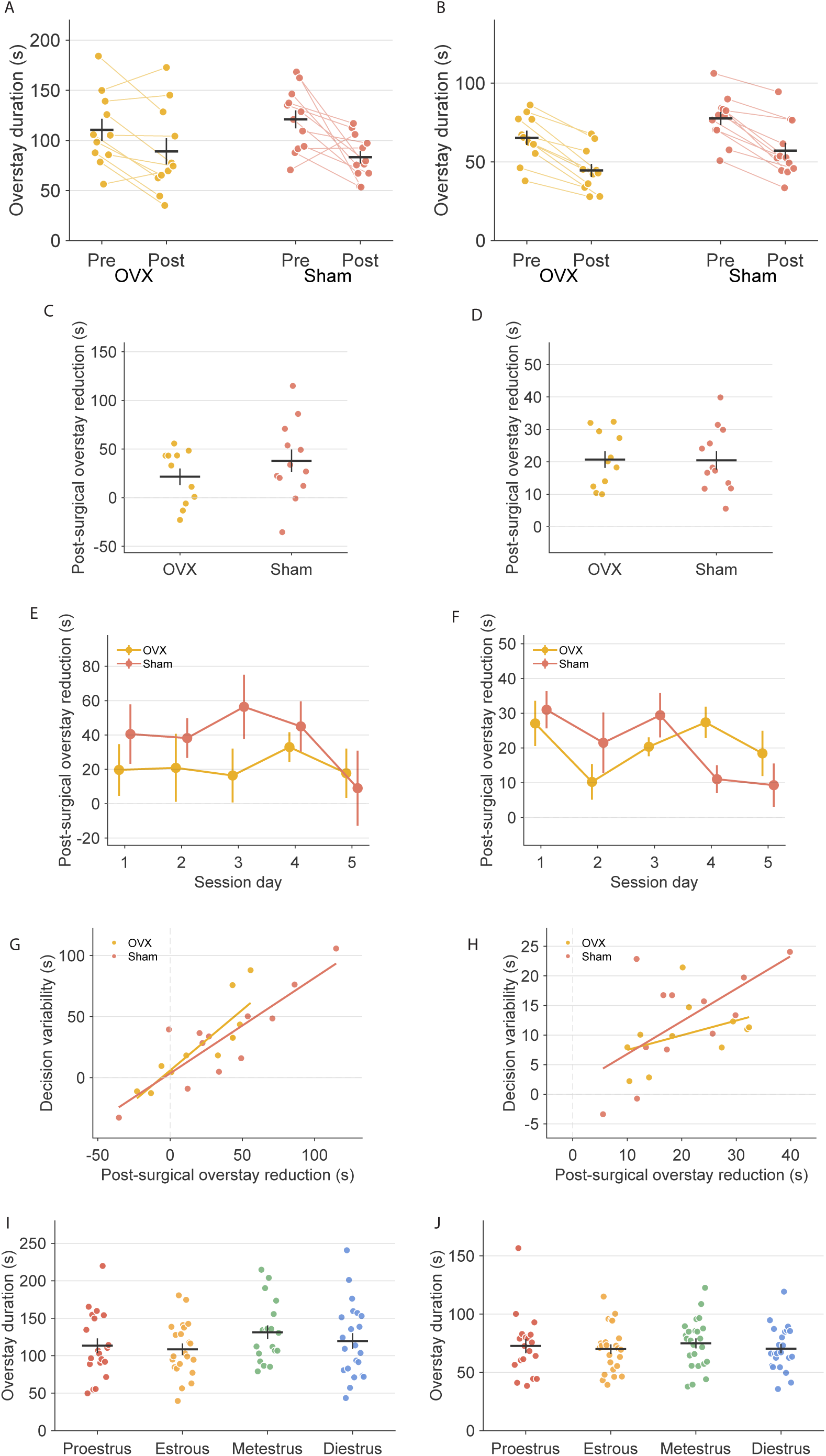
Female post-surgical overstay adaptation is modest and pre-surgical overstay is not clearly organized by estrous stage. A, pooled pre-surgical and post-surgical mean overstay duration in long-travel sessions for OVX and Sham females. B, pooled pre-surgical and post-surgical mean overstay duration in short-travel sessions. C, pooled long-travel post-surgical overstay reduction. D, pooled short-travel post-surgical overstay reduction. E, day-matched long-travel post-surgical overstay reduction across the five test sessions. F, day-matched short-travel post-surgical overstay reduction across the five test sessions. G, relationship between long-travel post-surgical overstay reduction and decision variability. H, relationship between short-travel post-surgical overstay reduction and decision variability. I, pre-surgical long-travel overstay grouped by estrous stage. J, pre-surgical short-travel overstay grouped by estrous stage. In panels A-D and G-J, each point represents one rat. In panels E and F, points and error bars indicate means ± SEM. For panels C-H, post-surgical overstay reduction was calculated as pre-surgical minus post-surgical overstay duration, such that larger positive values indicate a larger decrease in overstay after surgery. *p < 0.05; ns, not significant.

In short-travel sessions, both female groups showed lower post-surgical overstay than before surgery (Fig. 3B; ovariectomy: 65.28 ± 4.50 s pre, 44.58 ± 4.10 s post; sham: 77.56 ± 4.18 s pre, 57.10 ± 5.02 s post), but no significant post-surgical surgery × session interaction was detected (F(1,110) = 0.41, p = 0.5216). Consistent with this pattern, pooled post-surgical overstay reduction did not differ significantly between ovariectomized and sham females in either long-travel or short-travel sessions (Fig. 3C, D; long: p = 0.3401; short: p = 0.8294). Female overstay reduction was positively associated with overstay variability in both long-travel and short-travel conditions (Fig. 3G, H; long: rho = 0.792, p < 0.0001; short: rho = 0.508, p = 0.0144).

Because females underwent daily vaginal lavage throughout patch-leaving testing, we examined whether overstay varied across estrous stages and whether post-surgical cytology was consistent with successful ovariectomy. No significant estrous-stage effect was detected for pre-surgical long-travel overstay (Fig. 3I; one-way ANOVA, F(3,77) = 1.79, p = 0.1560) or pre-surgical short-travel overstay (Fig. 3J; F(3,84) = 0.53, p = 0.6656). Post-surgical vaginal cytology in ovariectomized females was consistent with loss of normal estrous cyclicity; therefore, post-surgical estrous-stage analyses were restricted to sham females. Within sham females, post-surgical overstay did not differ significantly across estrous stages in long-travel sessions (Supplementary Fig. 1A; F(3,41) = 1.04, p = 0.3833) or short-travel sessions (Supplementary Fig. 1B; F(3,42) = 0.41, p = 0.7474). Together, these findings indicate that the session-dependent long-travel effect observed in males was not mirrored after ovariectomy.

### Gonadectomy differentially alters spatial occupancy and idling in males and females

To determine whether sex and gonadectomy altered how rats allocated time within the patch, we combined males and females into a single rat-level occupancy framework and quantified stage-pooled patch-residence and idle-space occupancy ratios (Fig. 4A-D). Before surgery, both sexes concentrated occupancy in boundary and door-adjacent regions, but males showed a more center-biased distribution than females. In pre-surgical patch-residence occupancy, males showed lower door occupancy and higher center occupancy than females (Fig. 4A; door: males 16.9%, females 21.0%; Wilcoxon rank-sum test, p = 0.0172; center: males 23.8%, females 20.1%; p = 0.0091). In pre-surgical idle-space occupancy, males likewise showed higher idle-center occupancy than females (Fig. 4B; males 17.7%, females 14.3%; p = 0.0193), whereas females exhibited a higher overall idle-time ratio (27.3% vs 20.8%; p < 0.0001). After surgery, the overall occupancy structure remained qualitatively similar, but sex differences persisted in a narrower set of components. In post-surgical patch-residence occupancy, corridor occupancy remained lower in males than in females (Fig. 4C; males 4.5%, females 6.4%; p = 0.0018). In post-surgical idle-space occupancy, males showed lower corridor occupancy and higher center occupancy than females (Fig. 4D; corridor: males 5.9%, females 7.6%; p = 0.0280; center: males 19.8%, females 15.8%; p = 0.0480).

**Figure 4.**
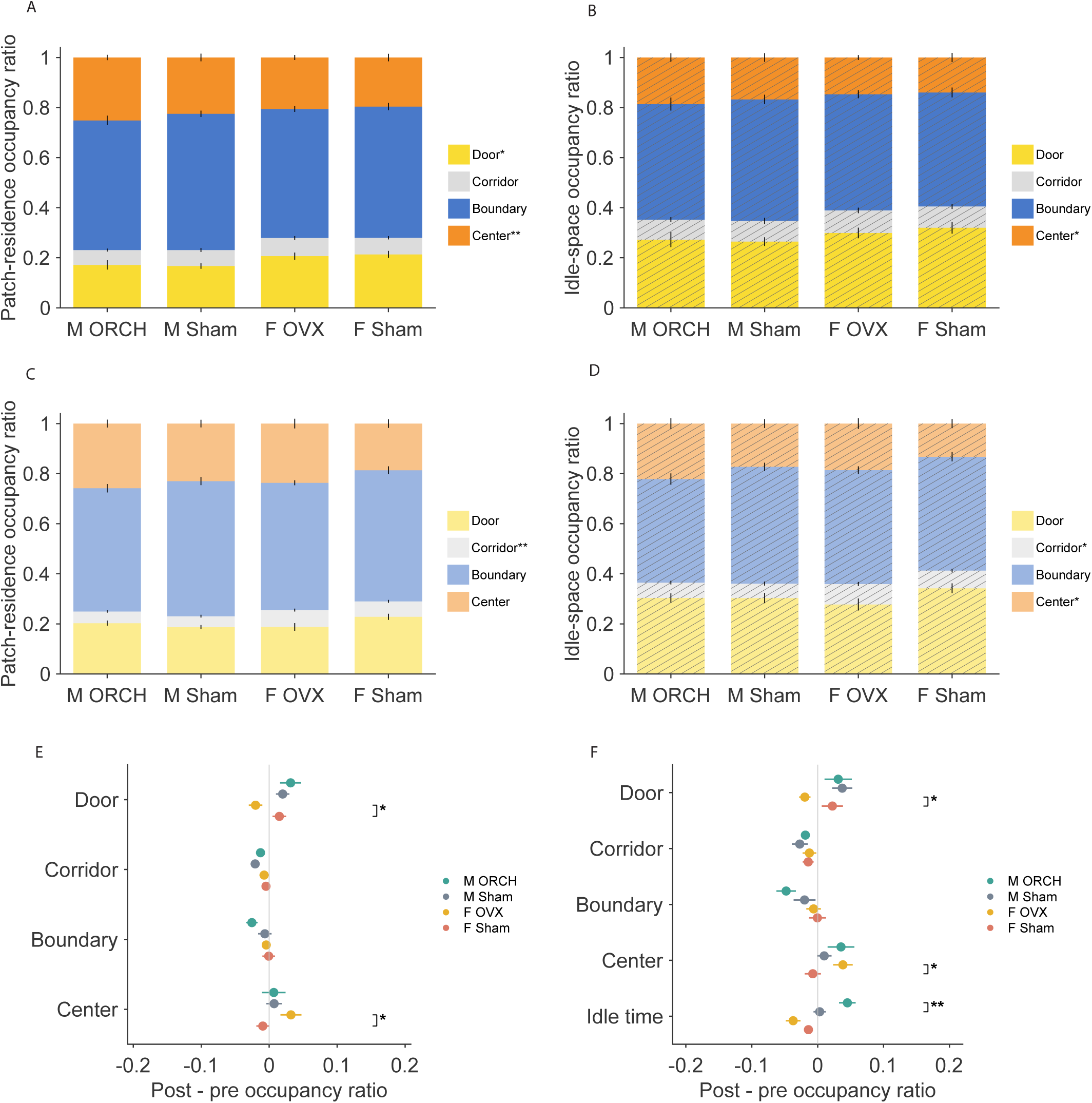
Patch-residence and idle-space occupancy show sex-specific spatial reorganization after gonadectomy. A, pre-surgical patch-residence occupancy composition for male ORCH, male Sham, female OVX, and female Sham groups pooled across long– and short-travel sessions. B, pre-surgical idle-space occupancy composition for the same four groups. C, post-surgical patch-residence occupancy composition. D, post-surgical idle-space occupancy composition. E, summary of post-pre change in patch-residence occupancy components across the four groups. F, summary of post-pre change in idle-space occupancy components, including idle-time ratio. In panels A-D, stacked bars show mean occupancy ratios for door, corridor, boundary, and center regions, and error bars denote SEM for the cumulative stacked components. Idle-space panels are displayed with diagonal hatching to distinguish them from patch-residence panels. In panels E and F, points and horizontal error bars indicate mean post-pre change ± SEM for each occupancy component. Positive values indicate an increase after surgery, and negative values indicate a decrease after surgery. *p < 0.05 for the within-sex surgery comparison for that occupancy component.

Post-pre change scores further showed that orchiectomy and ovariectomy altered occupancy in different ways (Fig. 4E, F). In males, patch-residence change was modest, but orchiectomy increased idle-time ratio relative to sham surgery (Fig. 4F; orchiectomy +4.5%, sham +0.3%; p = 0.0089). In females, ovariectomy produced a more spatially focused redistribution. OVX females showed a larger increase in patch center occupancy and a larger decrease in patch door occupancy than female shams (Fig. 4E; center: OVX +3.2%, sham –0.9%; p = 0.0337; door: OVX –2.0%, sham +1.5%; p = 0.0247). A parallel pattern was observed in idle-space change, where OVX females showed a larger increase in idle-center occupancy and a larger decrease in idle-door occupancy than female shams (Fig. 4F; center: OVX +3.8%, sham –0.7%; p = 0.0178; door: OVX –2.0%, sham +2.2%; p = 0.0455). When change scores were pooled by sex, males showed a larger increase in patch door occupancy than females (+2.6% vs –0.2%; p = 0.0187) and a markedly larger increase in idle-time ratio (+2.5% vs –2.5%; p < 0.0001). Together, these findings indicate that orchiectomy primarily increased post-surgical idling in males, whereas ovariectomy in females shifted occupancy away from the patch door and toward the patch center.

### Impulsive choice differed across surgical groups and showed a selective relationship with long-travel overstay reduction

Finally, because the impulsive-choice task was conducted after completion of post-surgical patch-leaving testing, we asked whether gonadal hormone status was associated with delay discounting and whether individual differences in impulsive choice were related to patch-leaving performance (Fig. 5A). Group differences in delayed reward choice were minimal at the shortest delays, with no significant effect at 0 s or 5 s (Fig. 5B; Kruskal-Wallis tests, p = 0.7925 and p = 0.3743, respectively). In contrast, significant group effects emerged at 15 s, 30 s, 50 s, and 75 s (p = 0.0163, p = 0.0006, p = 0.0019, and p = 0.0037, respectively). Across these longer delays, male sham rats maintained the strongest preference for the larger delayed reward, whereas orchiectomized males and both female groups showed steeper discounting.

**Figure 5.**
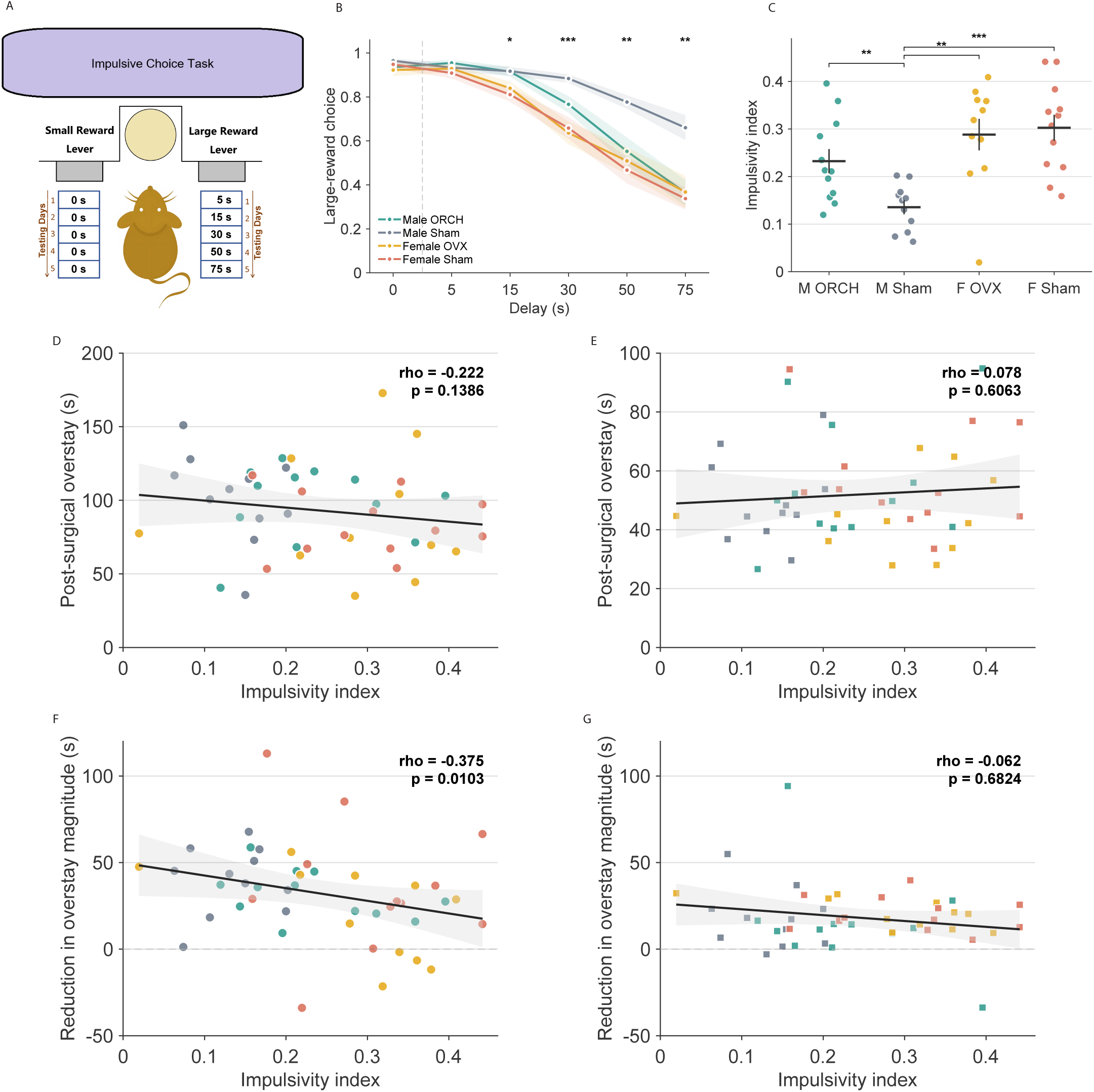
Gonadal status shapes impulsive choice and reveals an exploratory association with long-travel overstay reduction. A, schematic of the impulsive-choice task. B, delay-discounting curves for male ORCH, male Sham, female OVX, and female Sham rats. C, impulsivity index for each rat, with higher values indicating stronger delay discounting. D, relationship between impulsivity index and post-surgical long-travel overstay. E, relationship between impulsivity index and post-surgical short-travel overstay. F, relationship between impulsivity index and the reduction in long-travel overstay magnitude from pre– to post-surgery. G, relationship between impulsivity index and the corresponding short-travel reduction measure. In panels C-G, each point represents one rat. In panels D-G, lines denote least-squares fits for visualization, and inset text reports Spearman rho and p-values.

This group pattern was also reflected in the impulsivity index, for which higher values indicate greater impulsive choice (Fig. 5C). Male sham rats exhibited the lowest impulsivity index (0.136 ± 0.015), whereas orchiectomized males showed significantly higher impulsivity (0.232 ± 0.025; Wilcoxon rank-sum test, p = 0.0074). Male sham rats also showed significantly lower impulsivity than both OVX females (0.288 ± 0.033; p = 0.0013) and female sham rats (0.303 ± 0.027; p = 0.0003). Thus, removal of testicular hormones shifted males toward a more impulsive phenotype, whereas ovariectomy did not produce a comparably clear separation from female sham rats.

We next tested whether impulsive choice was related to post-surgical patch-leaving behavior across rats with both measures available (n = 46). Post-surgical overstay itself was not significantly associated with impulsivity in either long-travel sessions (Fig. 5D; Spearman rho = –0.222, p = 0.1386) or short-travel sessions (Fig. 5E; rho = 0.0780, p = 0.6063). By contrast, post-surgical overstay reduction in long-travel sessions was significantly associated with impulsive choice in the unadjusted analysis (Fig. 5F; rho = – 0.375, p = 0.0103), indicating that rats with higher impulsivity tended to show smaller reductions in overstay after surgery. No corresponding relationship was observed in short-travel sessions (Fig. 5G; rho = –0.0620, p = 0.6824). However, when sex and surgical group were included in the model, the association between impulsivity and long-travel overstay reduction was attenuated and no longer significant (linear model, beta = –81.36, p = 0.0809), suggesting that this relationship was partly accounted for by group structure. To summarize, these findings indicate that impulsive choice was related specifically to long-travel adjustment of patch-leaving behavior after surgery, rather than to post-surgical overstay magnitude itself.

## Discussion

The present study shows that gonadal hormone loss did not produce a uniform shift in patch-leaving behavior but instead revealed a selective role for testicular hormones in the adaptation of patch-leaving under high travel cost. Before surgery, males overstayed longer than females during long-travel sessions, whereas no clear sex difference was detected under short-travel conditions. This pattern is consistent with patch-foraging theory, which predicts that longer travel times increase the value of continued exploitation, and with recent rat work showing that sex differences in patch leaving become most apparent when travel cost is high ^1,2,4,10,11^. At the same time, both rodents and humans often remain in patches longer than simple reward-maximizing models predict, indicating that patch leaving reflects not only local depletion but also learning, uncertainty, opportunity-cost evaluation, and internal state ^4–12^. The present findings extend that literature by indicating that one component of this long-travel phenotype is hormonally regulated.

The clearest post-surgical effect emerged in males. Orchiectomy did not produce a global increase or decrease in overstay across all task conditions but instead attenuated the progressive reduction in long-travel overstay that remained evident across repeated post-surgical sessions in sham males. This result suggests that testicular hormones contribute less to a fixed leave threshold than to adaptive calibration of patch-leaving behavior when animals repeatedly encounter costly transitions. That interpretation is consistent with the broader view that patch leaving recruits prospective valuation and sensitivity to delayed future gains, computations that overlap with those engaged during intertemporal choice ^4,7,22,23^. It also fits evidence that androgens regulate mesocorticolimbic and prefrontal dopamine systems that support executive control, behavioral persistence, and cost-benefit evaluation ^15,28–32^. Notably, gonadectomy in male rats has been reported to reduce open-field activity, alter cortical catecholamine organization, modify prefrontal dopamine signaling, and impair performance on tasks that depend on prefrontal function ^28–32^. Together, these findings support the interpretation that the orchiectomy effect observed here reflects impaired adaptive adjustment under temporally costly foraging conditions rather than a nonspecific change in locomotion or motivation alone.

The female results were weaker and less directionally coherent. Although overstay generally declined after surgery in females, ovariectomized and sham females were not cleanly separated in the long-travel condition, and estrous stage did not significantly organize pre-surgical overstay. This pattern is broadly compatible with previous work showing that ovarian hormones can influence some forms of decision-making and behavioral allocation, including effort-based choice, memory-system bias, and risky decision-making, while effects on delay discounting itself are not always large or consistent across tasks ^13,14,16,17,20,33–35^. In that sense, the present data do not argue against ovarian hormone contributions to decision-making in general. Rather, they suggest that female patch-leaving overstay in this task was less strongly governed by ovarian status than male long-travel adaptation was by testicular status. The absence of a clear estrous-stage effect further suggests that natural cycling fluctuations in ovarian hormones were not a dominant source of variance in this specific patch-leaving measure, at least under the present sampling and task structure.

The occupancy analyses provided an important complement to the overstay data by showing that gonadectomy altered how animals behaved within the patch, not only when they left it. Before surgery, males spent relatively more time in patch-center regions and had lower idle-time ratios than females, indicating sex differences in the microstructure of patch engagement. After surgery, orchiectomy primarily increased idling in males, whereas ovariectomy more strongly redistributed female occupancy away from the patch door and toward the patch center. These dissociable patterns suggest that gonadal hormones influence patch-residence behavior through partially distinct mechanisms in the two sexes. In males, the increase in post-surgical idling is consistent with prior reports that gonadectomy reduces open-field activity and with evidence that testosterone modulates behavioral activation and persistence ^28,36^. In females, the OVX-related redistribution of occupancy may reflect altered exploratory allocation or altered weighting of spatial and motivational information within the patch rather than a strong shift in the terminal leave decision itself, which is consistent with literature showing estrogen-dependent modulation of exploratory and cognitive strategy use ^13,14,16,33,37^.

The impulsive-choice data place these foraging effects into a broader decision-making framework. Sham males showed the lowest impulsivity, whereas orchiectomized males shifted toward the female range, closely matching previous evidence that testicular hormones mediate robust sex differences in delay discounting in rats ^20^. More importantly, impulsivity was not clearly related to post-surgical overstay magnitude itself but was associated with long-travel overstay reduction in the unadjusted analysis. This pattern supports the idea that patch leaving and intertemporal choice share behaviorally relevant computations, specifically when leaving requires tolerance of delayed access to future benefits ^4,10,20,21,23^. However, that association was attenuated after accounting for sex and surgical group, indicating that the relationship is not well captured by a simple one-to-one trait correspondence across individuals. Instead, the shared signal may be partly expressed at the group level, with intact males combining lower impulsivity and stronger long-travel adaptation, whereas orchiectomized males shift in the opposite direction. This more restrained interpretation is also consistent with evidence that impulsive choice and impulsive action are dissociable constructs with overlapping but non-identical neural substrates ^25,38^.

Taken together, the results support a more specific conclusion than a general hormonal effect on patch leaving. The most robust phenotype in this dataset was a male-biased long-travel overstay pattern before surgery, followed by a selective loss of long-travel adaptation after orchiectomy, with weaker and less systematic effects after ovariectomy. In this sense, the major endocrine contribution identified here concerns adaptation to high travel cost rather than patch-leaving behavior in the abstract. This interpretation also helps reconcile the foraging and delay-discounting results: both point toward a role for testicular hormones in maintaining tolerance for delayed future outcomes, particularly when behavior must be adjusted across repeated experience rather than expressed as a fixed preference on a single session ^15,20,22,23,28^.

Several limitations should be considered. First, animals were singly housed, and the study did not include a sufficiently differentiated pre-surgical housing manipulation to evaluate whether housing-related stress interacted with gonadal status. This matters because social isolation can alter impulsive behavior in rats and may influence the overall behavioral range on which endocrine effects are expressed ^39,40^. Second, although the present design identifies behavioral consequences of gonadectomy, it does not establish which hormonal signals are sufficient to restore the observed phenotypes. Hormone-replacement experiments will therefore be important for determining whether testosterone replacement rescues long-travel adaptation and reduced idling in orchiectomized males, and whether estradiol replacement modifies the spatial-occupancy redistribution observed after ovariectomy ^14,15,20,29,30^. Third, the present task contrasted only short and long travel intervals. Adding an intermediate travel cost, such as 20 s, would help determine whether endocrine effects scale gradually with travel time or emerge only once delayed access to the next patch becomes sufficiently costly ^4,11^. Finally, future work should test whether the present long-travel phenotype is related not only to impulsive choice, as shown here, but also to impulsive action, which depends on partially dissociable control processes ^25,38,41^.

Overall, the present findings identify testicular hormones as a key contributor to sex differences in long-travel patch-leaving adaptation and show that gonadectomy reshapes both the timing and the microstructure of patch-residence behavior. More broadly, the data support the view that naturalistic foraging decisions and impulsive choice converge most strongly when animals must tolerate delayed access to future benefits, and that this convergence is especially sensitive to the presence of testicular hormones.

## Methods

### Subjects

All experimental procedures were approved by the Institutional Animal Care and Use Committee (IACUC) of the University of Illinois Urbana-Champaign. Forty-eight Long-Evans hooded rats (24 males and 24 females; approximately postnatal day 54 at arrival) were obtained from Charles River Laboratories in three cohorts. Animals were singly housed in a temperature-controlled room maintained at 23 degrees C under a 12 h light/dark cycle. During behavioral training and testing, body weight was maintained at >85% of free-feeding weight to promote food-motivated performance. Following surgery, one sham male (PLM90) and one ovariectomized female (PLF101) were excluded from post-surgical behavioral testing because they did not recover sufficiently to complete the remaining procedures.

### Patch-leaving apparatus

Patch-leaving behavior was tested in a custom acrylic apparatus composed of two square open fields (75 x 75 cm each) connected by a corridor (75 x 25 cm). The walls were 45 cm high, and extra-maze visual cues were placed around the apparatus to facilitate orientation. Access between the corridor and each open field was regulated by automated doors operated by stepper motors and triggered by infrared beam-break sensors. Above each open field, a 3D-printed sugar pellet dispenser delivered pellets into the arena at variable locations. All motors and sensors were controlled by an Arduino Uno board. A FLIR Flea3 camera mounted above the apparatus recorded behavior, and video and behavioral event streams were synchronized and acquired using Bonsai on a PC.

### Patch-leaving training and testing

Rats first underwent 5 days of handling and apparatus acclimation, with each handling session lasting 5 min. During acclimation, animals were placed in the corridor with both doors open, and 20 sugar pellets were distributed in each open field. Rats were then allowed to freely explore both patches for 10 min and consume the available pellets. Following acclimation, rats completed 5 days of patch-leaving training. During each training session, the rat was placed in the corridor with both doors closed. After 10 s, one door was opened pseudo randomly to allow entry into one patch and initiate the pellet-dispensing schedule in that open field. Rats were free to remain in the patch for any duration. When the rat exited the patch and approached the opposite door, triggering the infrared beam-break sensor, the door to the previous patch closed. After the programmed transit interval had elapsed, the opposite door opened and a new pellet-dispensing schedule began in the newly entered patch. This sequence could repeat throughout the 30-min session. Training was conducted using a short transit interval of 5 s. Patch-leaving testing was conducted individually and followed the same general procedure as training. Both the pre-surgical and post-surgical testing phases consisted of 10 daily sessions, with the first 5 sessions conducted under a long transit interval (60 s) and the final 5 sessions conducted under a short transit interval (5 s). The pellet-dispensing schedule was identical across training and testing. Specifically, pellet delivery within each patch followed a fixed depleting schedule in which the inter-pellet interval progressively increased over time. The same sequence was used in all sessions.

### Gonadectomy and sham surgeries

After completion of pre-surgical patch-leaving testing, rats underwent gonadectomy or sham surgery at approximately postnatal day 80. Surgeries were performed under isoflurane anesthesia (1.5-2.5%) with the animal placed on a sterile drape over a heating pad. Carprofen (5 mg/kg, s.c.) was administered immediately before surgery.

In females, bilateral ovariectomy was performed through small dorsal flank incisions made through the skin and muscle wall. Each ovary was exteriorized, ligated, and removed. In sham females, the same surgical exposure was performed without removing the ovaries. The muscle layer was closed with absorbable suture (Polyglactin 910), and the skin was closed with sterile silk sutures.

In males, orchiectomy was performed through a small ventral scrotal incision through the skin and cremaster muscle. Each testis was exteriorized and removed by severing the associated vasculature. In sham males, the same incision and surgical exposure were performed without removing the testes. The skin was closed with sterile silk sutures.

### Post-surgical care

Following surgery, rats were monitored during recovery from anesthesia until sternal recumbency and were then returned to the colony room. A second dose of carprofen (5 mg/kg, s.c.) was administered 6-24 h after the initial injection. Animals were monitored daily for 7 days for signs of pain, distress, dehydration, or impaired wound healing. Supportive care, including palatable food supplementation and warm 0.9% saline when needed, was provided during recovery. If required, additional carprofen was administered every 12-24 h until clinical signs improved. Sutures were removed within 10 days after surgery if they had not already detached spontaneously.

### Estrous cycle determination

The estrous stage was monitored during both the pre-surgical and post-surgical patch-leaving experiments by vaginal lavage. Lavage was performed daily at 9:00 AM throughout behavioral testing. Vaginal cytology samples were obtained by gently flushing the vaginal canal with room-temperature water using a P200 pipette and collecting the recovered fluid. Samples were placed onto standard glass microscope slides and stained with Toluidine blue. Cytological preparations were imaged under a brightfield microscope using a 10x objective. The estrous stage was identified from the relative abundance of nucleated epithelial cells, cornified epithelial cells, and leukocytes using standard criteria for Proestrus, Estrus, Metestrus, and Diestrus.

### Impulsive choice task

After completion of the patch-leaving experiment, rats remained food restricted at approximately 85% of their free-feeding body weight and were tested in an impulsive-choice task conducted in operant chambers using a delay-discounting procedure adapted from previously established methods ^39^. Lever assignments were counterbalanced across rats such that the smaller immediate reward and the larger delayed reward were assigned to opposite levers, and these assignments were held constant throughout training and testing.

Behavioral shaping proceeded in stages. Rats first completed 1 day of magazine training, during which 45 food pellets were delivered noncontingently at pseudorandom intervals ranging from 60 to 120 s while both levers remained retracted and the cue and house lights were off. Rats then underwent 6 days of lever-press training. During these sessions, only one lever was available at a time and the cue light above that lever was illuminated. Training alternated across days between the smaller-sooner and larger-later lever, with lever identity counterbalanced across animals and held constant thereafter. The first 2 training days included autoshaping, in which pellets were also delivered at variable intervals independent of responding.

If a rat earned fewer than 10 pellets during a session, supplemental hand-shaping was provided later the same day. Rats next completed 2 days of forced-choice training, in which each free-choice trial was followed by a trial in which only the previously unchosen lever was extended. After these shaping procedures, rats received 2 days of pre-training, followed by discrimination training with omission contingencies. During pre-training, both levers were available, reward delivery occurred 1 s after a lever press, and each session ended after 45 min or 45 trials, whichever came first. During discrimination training, failures to respond within 60 s were scored as omissions and advanced directly to the inter-trial interval. Rats proceeded to testing after choosing the larger reward on at least 80% of trials for 3 consecutive days.

Testing was carried out across 5 consecutive daily sessions and was identical to discrimination training except that the larger reward was delivered after an imposed delay. The smaller-sooner option always delivered 1 pellet after 1 s, whereas the larger-later option delivered 3 pellets after a fixed delay that increased across test days (5, 15, 30, 50, and 75 s). Session structure remained otherwise unchanged, including omission trials, and rats advanced through the delay series regardless of performance on the previous day. Impulsive choice was quantified as the preference ratio for the larger-later option at each delay, calculated as the proportion of completed trials on which the larger-reward lever was selected. Higher ratios therefore indicate a greater preference for the delayed larger reward and lower impulsive choice. For summary analyses, discounting across delays was additionally reduced to a normalized area-under-the-curve measure, and the impulsivity index was defined as 1 – AUC, with larger values indicating greater impulsive choice.

### Behavioral measures

Patch-leaving performance was quantified from the trial-wise stay durations recorded in each session. Unless otherwise specified, session-level values were averaged within each rat across the relevant five-day block for a given travel condition and surgical stage. To quantify surgical change in patch-leaving behavior, post-surgical overstay reduction was defined as pre-surgical mean overstay minus post-surgical mean overstay, such that larger positive values indicate a greater reduction in overstay after surgery. For day-matched analyses, this calculation was performed separately for corresponding pre– and post-surgical sessions (days 1-5). When overstay magnitude was analyzed without regard to sign, absolute overstay reduction was defined analogously using absolute overstay duration.

Decision variability was used to quantify trial-to-trial consistency in patch-leaving behavior within a session. For each session, decision variability was calculated as the standard deviation of trial-wise overstay values, such that lower values reflected more stable patch-leaving decisions. Post-surgical reduction in decision variability was defined as pre-surgical variability minus post-surgical variability, so that positive values indicate increased decision stability after surgery.

Locomotor and spatial measures were derived from x-y position tracking sampled at 30 frames/s. Speed was calculated from frame-to-frame displacement and converted to cm/s using a tracking calibration factor of 6.5 pixels per cm. Frames with speed < 3 cm/s were classified as idle. Idle time ratio was calculated as the number of idle frames divided by the total number of patch-residence frames. Patch-residence spatial occupancy was calculated as the proportion of patch-residence frames spent within each predefined region of the apparatus. Idle occupancy was calculated as the proportion of idle frames spent within each region. For occupancy change analyses, post-surgical change was defined as post-surgical minus pre-surgical values.

Impulsive choice was quantified from the delay-discounting task using a normalized area-under-the-curve approach. For each rat, preference for the larger delayed reward was calculated at each delay as the proportion of completed trials on which the larger reward was chosen. The area under the delay-discounting curve (AUC) was then calculated across ordered delay bins, and the impulsivity index was defined as 1 – AUC, such that larger values indicated steeper discounting and greater impulsive choice.

## Statistical analysis

All analyses were conducted in MATLAB 2022b using custom scripts. The rat was treated as the unit of analysis throughout. Unless otherwise specified, session-level values were first summarized within rat, and data are presented as mean ± SEM.

Because sample sizes were modest and several pooled behavioral measures were not assumed to be normally distributed, independent-group comparisons were generally performed using two-sided Wilcoxon rank-sum tests, and paired pre-versus post-surgical comparisons within the same rats were performed using two-sided Wilcoxon signed-rank tests. These tests were used for baseline sex comparisons, pooled sham-versus-gonadectomy comparisons within sex, and change-score analyses. Delay-discounting curves were compared across groups at each delay bin using Kruskal-Wallis tests, and planned pairwise comparisons of impulsivity index between groups were performed using Wilcoxon rank-sum tests. Associations between behavioral measures were first assessed using Spearman rank correlations.

To examine day-by-day changes in patch-leaving behavior across the five testing sessions, linear mixed-effects models were fit to session-level overstay data with rat identity included as a random intercept to account for repeated observations within animals. Fixed effects included surgical group, surgical stage, session day, and the relevant sex term or sex-specific subgrouping for the comparison of interest. Session day was modeled to test whether surgery altered the trajectory of across-session adaptation rather than only the pooled pre-versus post-surgical mean. Follow-up models were conducted separately for long-and short-travel conditions and, where appropriate, separately within each sex. To directly compare male and female post-surgical long-travel trajectories, a combined model including sex, surgical group, and session day interaction terms was used.

For impulsivity analyses, significant bivariate associations were further evaluated using follow-up linear models that included sex and surgical group as covariates to determine whether the observed relationship remained after accounting for group structure.

Estrous-cycle effects on pre-surgical female overstay were assessed using one-way ANOVA across estrous stages. All tests were two-sided, and p < 0.05 was considered statistically significant.

## Acknowledgements

We thank PoChing Patrick Lin for guidance and surgical instruction in the orchiectomy procedure and Amara Brinks for guidance and instruction in the ovariectomy procedure.

## Author contributions

Y.D. and J.R.H. conceived and designed the study. Y.D. and K.C. collected the data. Y.D. analyzed the data, prepared the figures, and drafted the manuscript. J.R.H. supervised the study and edited the manuscript. All authors reviewed and approved the final manuscript.

## Data availability

The datasets generated during and/or analyzed during the current study are available from the corresponding author on reasonable request.

## FIGURE LEGENDS

**Supplementary Figure 1.**
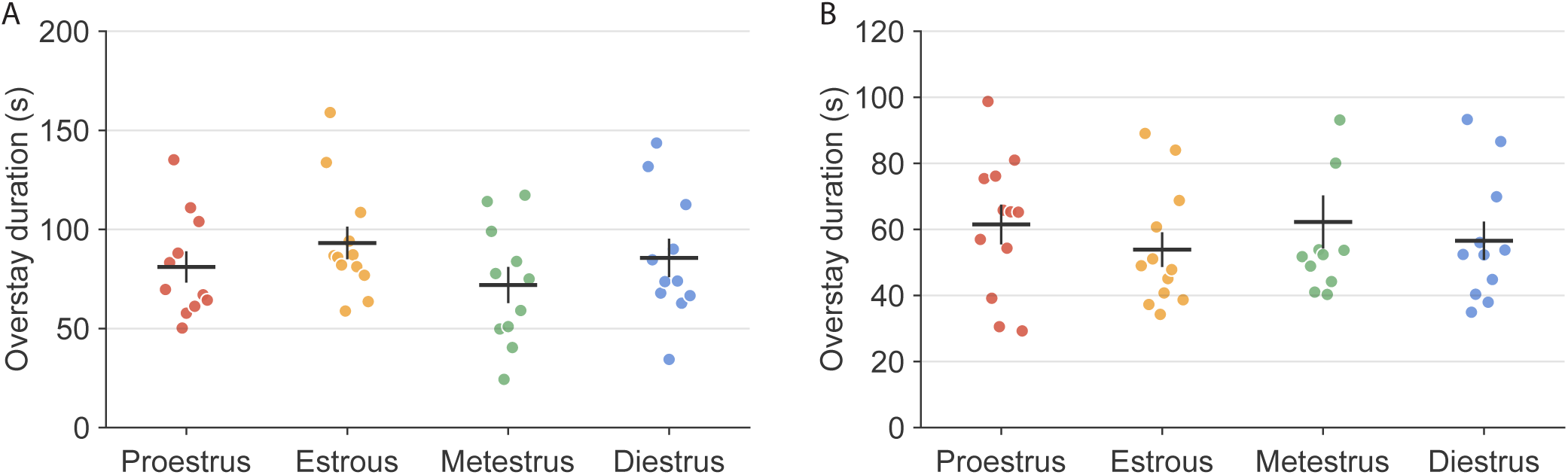
Post-surgical overstay in sham females remains not clearly organized by estrous phase. A, post-surgical long-travel overstay in sham females grouped by lavage-defined estrous phase. B, post-surgical short-travel overstay in sham females grouped by lavage-defined estrous phase. In both panels, each point represents one rat averaged across all post-surgical sessions observed in that cycle state; horizontal bars denote means and vertical bars denote SEM.

## References

1. MacArthur, R. H. & Pianka, E. R. On Optimal Use of a Patchy Environment. The American Naturalist 100, 603–609 (1966).

2. Charnov, E. L. Optimal foraging, the marginal value theorem. Theoretical Population Biology 9, 129–136 (1976).

3. Stephens, D. W. Decision ecology: Foraging and the ecology of animal decision making. *Cognitive, Affective*, & Behavioral Neuroscience 8, 475–484 (2008).

4. Kane, G. A. et al. Rats exhibit similar biases in foraging and intertemporal choice tasks. eLife 8, e48429 (2019).

5. Kane, G. A. et al. Rat Anterior Cingulate Cortex Continuously Signals Decision Variables in a Patch Foraging Task. J. Neurosci. 42, 5730–5744 (2022).

6. Gancarz, A. M. et al. Reward maximization assessed using a sequential patch depletion task in a large sample of heterogeneous stock rats. Sci Rep 13, 7027 (2023).

7. Hayden, B. Y., Pearson, J. M. & Platt, M. L. Neuronal basis of sequential foraging decisions in a patchy environment. Nat Neurosci 14, 933–939 (2011).

8. López-Yépez, J. S., Martin, J., Hulme, O. & Kvitsiani, D. Choice history effects in mice and humans improve reward harvesting efficiency. PLoS Comput Biol 17, e1009452 (2021).

9. Harhen, N. C. & Bornstein, A. M. Overharvesting in human patch foraging reflects rational structure learning and adaptive planning. Proc. Natl. Acad. Sci. U.S.A. 120, e2216524120 (2023).

10. Dai, Y., Seielstad, A. K. L. & Hinman, J. R. Social isolation biases female rats toward safety-oriented, efficient foraging. bioRxiv 2026.07.03.736107 (2026) doi:10.64898/2026.07.03.736107.

11. Garcia, M., Gupta, S. & Wikenheiser, A. M. Sex differences in patch-leaving foraging decisions in rats. Oxford Open Neuroscience 2, kvad011 (2023).

12. Kane, G. A. et al. Increased locus coeruleus tonic activity causes disengagement from a patch-foraging task. Cogn Affect Behav Neurosci 17, 1073–1083 (2017).

13. Frick, K. M., Kim, J., Tuscher, J. J. & Fortress, A. M. Sex steroid hormones matter for learning and memory: estrogenic regulation of hippocampal function in male and female rodents. Learn. Mem. 22, 472–493 (2015).

14. Uban, K. A., Rummel, J., Floresco, S. B. & Galea, L. A. M. Estradiol Modulates Effort-Based Decision Making in Female Rats. Neuropsychopharmacol 37, 390–401 (2012).

15. Tobiansky, D. J., Wallin-Miller, K. G., Floresco, S. B., Wood, R. I. & Soma, K. K. Androgen Regulation of the Mesocorticolimbic System and Executive Function. Front. Endocrinol. 9, 279 (2018).

16. Korol, D. L. Role of estrogen in balancing contributions from multiple memory systems. Neurobiology of Learning and Memory 82, 309–323 (2004).

17. Becker, J. B. & Koob, G. F. Sex Differences in Animal Models: Focus on Addiction. Pharmacological Reviews 68, 242–263 (2016).

18. Larson, E. B. & Carroll, M. E. Estrogen Receptor β, but not α, Mediates Estrogen’s Effect on Cocaine-Induced Reinstatement of Extinguished Cocaine-Seeking Behavior in Ovariectomized Female Rats. Neuropsychopharmacol 32, 1334–1345 (2007).

19. Wood, R. I. et al. ’Roid rage in rats? Testosterone effects on aggressive motivation, impulsivity and tyrosine hydroxylase. Physiology & behavior 110-111, 6–12 (2013).

20. Hernandez, C. M. et al. Testicular hormones mediate robust sex differences in impulsive choice in rats. eLife 9, e58604 (2020).

21. Evenden, J. L. & Ryan, C. N. The pharmacology of impulsive behaviour in rats: the effects of drugs on response choice with varying delays of reinforcement. Psychopharmacology 128, 161–170 (1996).

22. Floresco, S. B., Onge, J. R. St., Ghods-Sharifi, S. & Winstanley, C. A. Cortico-limbic-striatal circuits subserving different forms of cost-benefit decision making. Cognitive, Affective, & Behavioral Neuroscience 8, 375–389 (2008).

23. Frost, R. & McNaughton, N. The neural basis of delay discounting: A review and preliminary model. Neuroscience & Biobehavioral Reviews 79, 48–65 (2017).

24. Cross, C. P., Copping, L. T. & Campbell, A. Sex differences in impulsivity: a meta-analysis. Psychol Bull 137, 97–130 (2011).

25. Weafer, J. & De Wit, H. Sex differences in impulsive action and impulsive choice. Addictive Behaviors 39, 1573–1579 (2014).

26. Eubig, P. A., Noe, T. E., Floresco, S. B., Sable, J. J. & Schantz, S. L. Sex differences in response to amphetamine in adult Long–Evans rats performing a delay-discounting task. Pharmacology Biochemistry and Behavior 118, 1–9 (2014).

27. Garcia, A. & Kirkpatrick, K. Impulsive choice behavior in four strains of rats: Evaluation of possible models of Attention-Deficit/Hyperactivity Disorder. Behavioural Brain Research 238, 10–22 (2013).

28. Adler, A., Vescovo, P., Robinson, J. K. & Kritzer, M. F. Gonadectomy in adult life increases tyrosine hydroxylase immunoreactivity in the prefrontal cortex and decreases open field activity in male rats. Neuroscience 89, 939–954 (1999).

29. Aubele, T. & Kritzer, M. F. Androgen Influence on Prefrontal Dopamine Systems in Adult Male Rats: Localization of Cognate Intracellular Receptors in Medial Prefrontal Projections to the Ventral Tegmental Area and Effects of Gonadectomy and Hormone Replacement on Glutamate-Stimulated Extracellular Dopamine Level. Cerebral Cortex 22, 1799–1812 (2012).

30. Aubele, T. & Kritzer, M. F. Gonadectomy and Hormone Replacement Affects In Vivo Basal Extracellular Dopamine Levels in the Prefrontal Cortex but Not Motor Cortex of Adult Male Rats. Cerebral Cortex 21, 222–232 (2011).

31. Kritzer, M. F., Brewer, A., Montalmant, F., Davenport, M. & Robinson, J. K. Effects of gonadectomy on performance in operant tasks measuring prefrontal cortical function in adult male rats. Hormones and Behavior 51, 183–194 (2007).

32. Kritzer, M. F. Regionally Selective Effects of Gonadectomy on Cortical Catecholamine Innervation in Adult Male Rats Are Most Disruptive to Afferents in Prefrontal Cortex. Cerebral Cortex 9, 507–518 (1999).

33. Westbrook, S. R., Hankosky, E. R., Dwyer, M. R. & Gulley, J. M. Age and sex differences in behavioral flexibility, sensitivity to reward value, and risky decision-making. Behavioral Neuroscience 132, 75–87 (2018).

34. Orsini, C. A., Willis, M. L., Gilbert, R. J., Bizon, J. L. & Setlow, B. Sex differences in a rat model of risky decision making. Behavioral Neuroscience 130, 50–61 (2016).

35. Orsini, C. A. et al. Regulation of risky decision making by gonadal hormones in males and females. Neuropsychopharmacol. 46, 603–613 (2021).

36. Justel, N., Ruetti, E., Bentosela, M., Mustaca, A. E. & Papini, M. R. Effects of testosterone administration and gonadectomy on incentive downshift and open field activity in rats. Physiology & Behavior 106, 657–663 (2012).

37. Bronstein, P. M. & Hirsch, S. M. Reactivity in the rat: Ovariectomy fails to affect open-field behaviors. Bull. Psychon. Soc. 3, 257–260 (1974).

38. Wang, Q. et al. Dissociated neural substrates underlying impulsive choice and impulsive action. NeuroImage 134, 540–549 (2016).

39. Venegas, J. J., et al. Social isolation increases impulsive choice with minor changes on metabolic function in middle-aged rats. Physiological Reports 13, e70184 (2025).

40. Lovic, V., Keen, D., Fletcher, P. J. & Fleming, A. S. Early-life maternal separation and social isolation produce an increase in impulsive action but not impulsive choice. Behavioral Neuroscience 125, 481–491 (2011).

41. Beatty, W. W. Effects of gonadectomy on sex differences in DRL behavior. Physiology & Behavior 10, 177–178 (1973).

